# TvAP65 in *Trichomonas vaginalis* Promotes HPV Infection by Interacting with Host Molecules

**DOI:** 10.1101/2024.09.27.615334

**Authors:** Xuefang Mei, Wanxin Sheng, Yani Zhang, Wenjie Tian, Xiaowei Tian, Zhenke Yang, Shuai Wang, Zhenchao Zhang

## Abstract

Cervical cancer induced by human papillomavirus (HPV) infection poses a serious threat to women’s health. Studies have shown that *Trichomonas vaginalis* (*T. vaginalis*), which is widely prevalent globally, can facilitate HPV infection. However, the underlying mechanism remains unclear. This study found that *T. vaginalis* significantly enhanced HPV infection in HaCaT cells and mouse vaginas through *in vivo* and *in vitro* experiments, and promoted the expression of HPV membrane receptor molecules CD151 and HSPG2. The HPV infection rate and CD151/HSPG2 expression levels were significantly decreased after reducing the expression of *T. vaginalis* adhesion protein 65 (TvAP65). In contrast, both HPV infection rates and CD151/HSPG2 expression were significantly increased in HaCaT cells over-expressing TvAP65. When both TvAP65 in *T. vaginalis* and CD151/HSPG2 in HaCaT cells were knocked down simultaneously, the infection rate of HPV in HaCaT cells was further reduced. These results suggest that TvAP65 promotes HPV infection by up-regulating the expression of CD151 and HSPG2. Furthermore, this study knocked down the 12 interacting molecules of TvAP65 in HaCaT cells one by one, and found that the HPV infection rate was significantly reduced after *T. vaginalis* infected HaCaT cells with low expression of FTH1, SPCS1, ATP5MC3, ITGB7, PMEPA1 or REEP5. Among them, SPCS1 played the most significant role. Simultaneous knockdown of TvAP65 and SPCS1 further significantly down-regulated the infection rate of HPV in HaCaT cells. Moreover, this molecule also down-regulated the promoting effect of *T. vaginalis* on HSPG2/CD151 expression. These results imply that SPCS1 plays an important role in *T. vaginalis* promoting HSPG2/CD151 expression and HPV infection. This study not only further proved that *T. vaginalis* can promote HPV infection but also explores the molecular mechanism by which TvAP65, through its interaction with SPCS1, up-regulates the expression of HSPG2 and CD151, thereby facilitating HPV infection. This provides a theoretical basis for clarifying the mechanism of co-infection between *T. vaginalis* and HPV.

## Background

*Trichomonas vaginalis* (*T. vaginalis)* is one of the most common non-viral sexually transmitted pathogens causing reproductive tract infections [1]. It can establish persistent infections in the vagina. Approximately 276 million people worldwide are infected with *T. vaginalis* each year [2]. Due to the large number of asymptomatic patients, the number of undiagnosed cases is quite high [3]. Nevertheless, the preferred diagnostic methods in many regions have low sensitivity, leading to a high rate of missed diagnoses [4]. Additionally, because trichomoniasis is not a notifiable disease, these epidemiological data may be underestimated [5]. Consequently, the Centres for Disease Control and Prevention (CDC) has listed trichomoniasis as one of the neglected parasitic infections (NPIs)[6].

Infection with *T. vaginalis* can cause a number of complications, including vaginitis, urethritis, cervicitis, prostatitis, and increased risk of acquisition and transmission of the human immunodeficiency virus, and cancer [7, 8, 9]. Pregnant women infected with *T. vaginalis* are more likely to have low birth weight infants and preterm births [10]. Although trichomoniasis is considered a relatively milder and curable disease, it has a high incidence, and a growing number of studies are showing resistance to metronidazole [11, 12]. In addition, epidemiological studies have shown that *T. vaginalis* infection leads to an increased risk of cervical cancer [13, 14, 15, 16]. However, the mechanism by which *T. vaginalis* causes cervical cancer has not been fully elucidated. It has been reported that the inflammatory process caused by *T. vaginalis* destroys cervical epithelial cells, which promotes the entry of HPV into the basal layer of the cervical epithelium [17, 18, 19, 20]. Consequently, this leads to the integration of viral DNA into host DNA and to the overexpression of viral oncogenes, thereby activating the carcinogenic mechanisms [18, 20, 21].

Cervical cancer is a major global health problem. The incidence of cervical cancer exceeds 528,000 cases annually, with more than 270,000 deaths [22]. High-risk HPV infection is the primary cause of cervical cancer, with more than 70% of cervical cancers caused by HPV16 and HPV18 infections [23]. This disease progresses rapidly and has a high mortality rate, posing a severe threat to women’s health [24]. Studies have shown that infections with HPV16, 31, and 33 subtypes are associated with the clinical course of trichomoniasis [25, 26]. The risk of HPV16 infection increases 6.5 times in the presence of *T. vaginalis* [13]. A meta-analysis on the relationship between *T. vaginalis* and HPV infection found that the HPV infection rate in *T. vaginalis*-positive patients was significantly higher than in normal women [27], further confirming that *T. vaginalis* is a risk factor for HPV infection. Additionally, there is evidence of a higher cancer risk observed in women with both HPV and *T. vaginalis* infection [19]. Despite the significant effectiveness of prophylactic vaccines against common HPV subtypes, existing vaccines cannot prevent all pathogenic HPV subtypes, and vaccine coverage is limited in low- and middle-income countries [28]. Therefore, the impact of *T. vaginalis* on HPV infection remains an important public health issue.

As an extracellular parasite, the adhesion of *T. vaginalis* to host cells is a prerequisite for its parasitism and pathogenicity [29]. This process is mediated by five adhesion proteins on the surface of *T. vaginalis*, including TvAP23, TvAP33, TvAP51, TvAP65, and TvAP120. Among these, TvAP65 plays a significant role in mediating the binding of the parasite to host cells [30] and is essential in the adhesion to host cells, inhibition of host cell proliferation, induction of host cell apoptosis, and death [31]. Additionally, passive immunisation with anti-rTvAP65 PcAbs or blocking TvAP65 protein can reduce the pathogenicity of *T. vaginalis* [31]. This study aims to systematically investigate the role and mechanism of TvAP65 of *T. vaginalis* in HPV infection at the molecular, cellular, and animal levels, providing a basis for elucidating the mechanism by which *T. vaginalis* promotes HPV infection.

## Materials and methods

### Ethical statement

All experiments involved in this study were reviewed and approved by the Ethics Review Committee of Xinxiang Medical College (Reference No. XYLL-20210184). The 6-week-old female BALB/cA-nu mice of SPF grade were fed sterile feed and purified water in an animal house with appropriate humidity and temperature. When the infected mice were near death, they were humanely euthanized by exposure to 60-70% carbon dioxide for 5 minutes, making every effort to minimize their suffering. Occasionally, cervical dislocation was used to confirm the effectiveness of euthanasia.

### Cell and *T. vaginalis* culture

DMEM medium (Procell Life Science & Technology Co., Ltd., Wuhan, China) containing 10% foetal bovine serum (FBS; Procell Life Science & Technology Co., Ltd.) and 1% penicillin-streptomycin (Procell Life Science & Technology Co., Ltd.) was used for HaCaT cell culture. Cells were evenly plated in cell culture plates and placed in a cell incubator containing 5% CO_2_ at 37℃. The culture medium was changed according to cell growth status. When cells reached the logarithmic growth phase, 0.25% trypsin was added to cover the cell surface for digestion, incubated at 37°C for 4∼5 min, then terminated with the addition of medium and centrifuged at 1000 rpm for 3 min. Cells were then passaged in new culture plates for passage culture.

TYM medium (TUOPU Biol-engineering Co., Ltd., Shandong, China) containing 20% FBS (Procell Life Science & Technology Co., Ltd.) and 1% penicillin-streptomycin (Procell Life Science & Technology Co., Ltd.) was used for *T. vaginalis* culture. Parasites were suspended in test tubes and cultured in a cell incubator containing 5% CO_2_ at 37℃. When *T. vaginalis* reached the logarithmic growth phase, they were inoculated at a 1: 50 ratio in new TYM medium for subculture.

### RNA interference

RNA interference (RNAi) technology was used to reduce gene expression levels in this study. Among them, the siRNA of CD151, HSPG2 and their corresponding negative control were purchased from Santa Cruz Biotechnology Co., Ltd., Shanghai, China, and the product code were sc-42829, sc-44010 and sc-37007, respectively. The siRNA sequences of TvAP65 and its interaction molecules were designed and synthesized by Suzhou Jinweizhi Biotechnology Co., Ltd., Suzhou, China. Three pairs of siRNAs were synthesized for each gene and one pair with the highest interference efficiency was selected. The siRNA sequences are listed in Table 1.

**Table 1.**
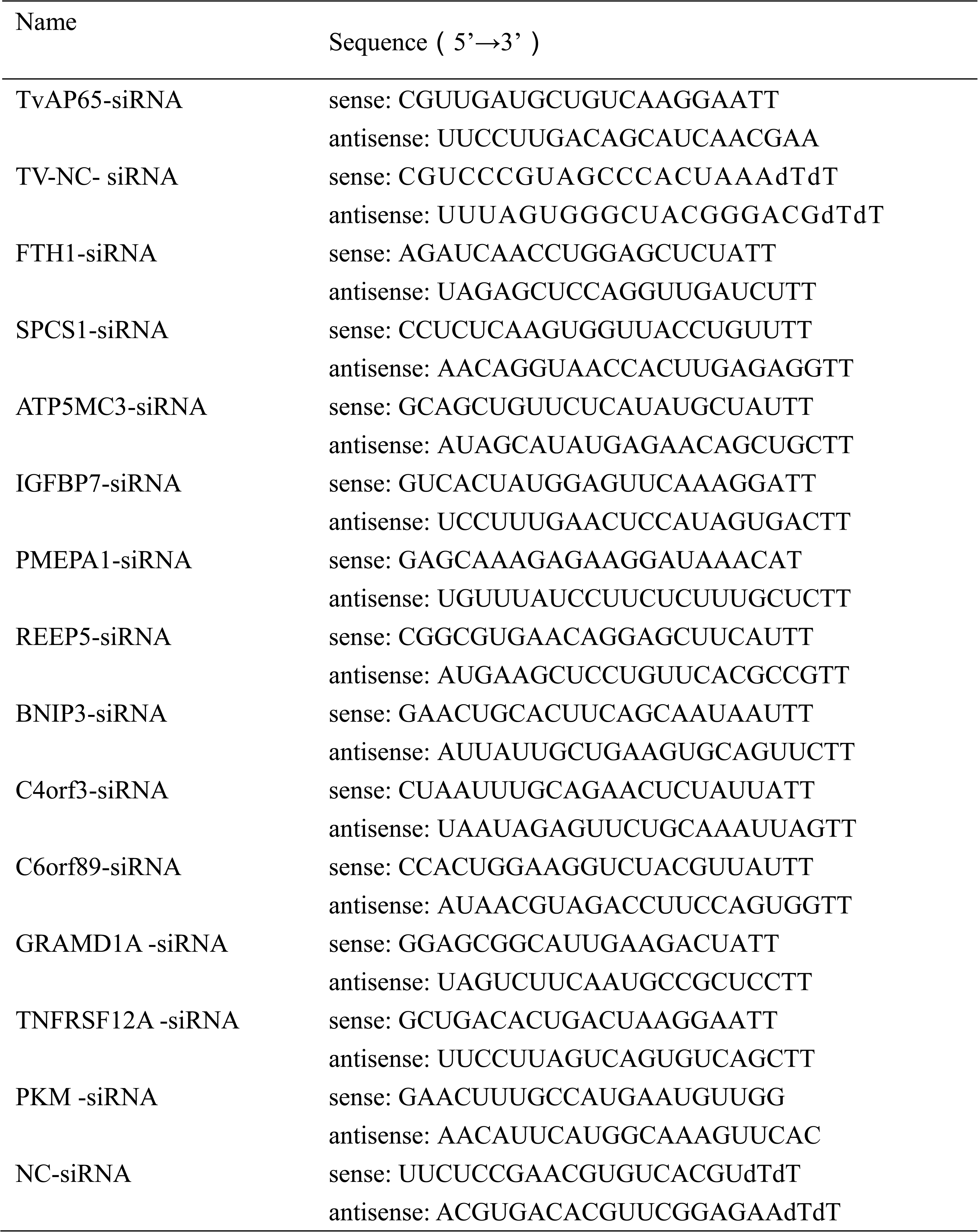
The siRNA sequences of TvAP65 and its interacting molecules.

For TvAP65 gene of *T. vaginalis*, 5×10^5^ *T. vaginalis* were resuspended in 400 μL serum-free, antibiotic-free TYM basal medium (TUOPU Biol-engineering Co., Ltd.) and cultured in a 24-well cell culture plate. 5 μL of 20 μM TvAP65-siRNA was mixed with 50 μL opti-MEM I reduced-serum medium (Thermo Fisher Scientific, Waltham, MA, USA) and gently pipetted. 1.2 μL Lipofectamine 2000 (Thermo Fisher Scientific) was mixed with 50 μL opti-MEM I reduced-serum medium (Thermo Fisher Scientific) and incubated at room temperature for 5 min. The two mixtures were then combined and incubated at room temperature in the dark for 20 min to prepare the transfection reagent complex. The transfection reagent complex was added to 5×10^5^ *T. vaginalis* and cultured in a cell incubator containing 5% CO_2_ at 37℃. After 12 h, 500 μL complete TYM medium (TUOPU Biol-engineering Co., Ltd.) was added and cultured for another 12 h. Simultaneously, a blank control group, a Lipofectamine 2000 group, and an NC-siRNA group were set up, with three replicates per group. *T. vaginalis* from each group were collected, and TvAP65 gene expression was detected by qPCR (primer sequence is shown in Table 2) and western blot (see additional file 1: Figure S1).

**Table 2.**
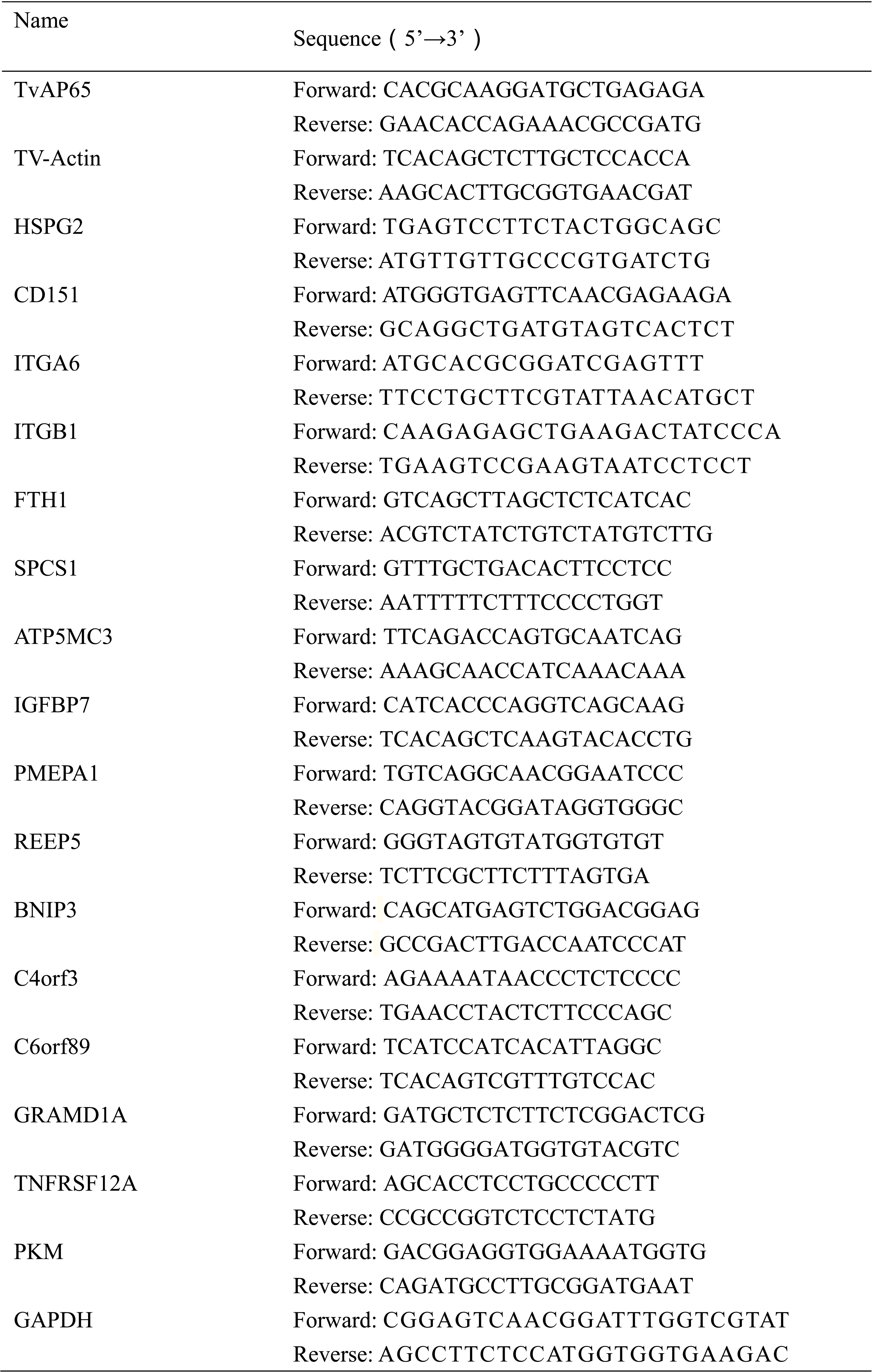
Sequences of primers used for qPCR.

For HSPG2, CD151, FTH1, SPCS1, ATP5MC3, IGFBP7, PMEPA1, REEP5, BNIP3, C4orf3, C6orf89, GRAMD1A, TNFRSF12A, and PKM gene in HaCaT cells, 1 × 10^6^ HaCaT cells were resuspended in 1.5 mL opti-MEM I reduced-serum medium (Thermo Fisher Scientific) and cultured in a 6-well cell culture plate. 10 μL of 10 μM siRNA for CD151 or HSPG2 genes, or 20 μM siRNA for FTH1, SPCS1, ATP5MC3, IGFBP7, PMEPA1, REEP5, BNIP3, C4orf3, C6orf89, GRAMD1A, TNFRSF12A, or PKM genes were mixed with 250 μL opti-MEM I reduced-serum medium (Thermo Fisher Scientific) and gently pipetted. 5 μL Lipofectamine 2000 (Thermo Fisher Scientific) was mixed with 250 μL opti-MEM I reduced-serum medium (Thermo Fisher Scientific) and incubated at room temperature for 5 min. The two mixtures were then combined and incubated at room temperature in the dark for 20 min to prepare the transfection reagent complex. The transfection reagent complex was added to 1×10^6^ HaCaT cells and cultured in a cell incubator containing 5% CO_2_ at 37℃. Simultaneously, a blank control group, a Lipofectamine 2000 group, and an NC-siRNA group were set up, with three replicates per group. HaCaT cells from each group were collected, and gene expression was detected by qPCR (primer sequence is shown in Table 2) and western blot (see additional file 2: Figure S2 and additional file 3: Figure S3).

### The transfection of pDsRed-N1-TvAP65 eukaryotic expression vector into HaCaT cells

1×10^6^ HaCaT cells were re-suspended in 1.5 mL opti-MEM I reduced-serum medium (Thermo Fisher Scientific) and cultured in a 6-well cell culture plate. 2500 ng pDsRed-N1-TvAP65 eukaryotic expression vector was re-suspended in 250 μL opti-MEM I reduced-serum medium (Thermo Fisher Scientific), gently pipetted, and incubated at room temperature for 5 min in the dark. 10 μL of Lipofectamine 2000 (Thermo Fisher Scientific) was mixed with 250 μL of opti-MEM I reduced-serum medium (Thermo Fisher Scientific) and. The two mixtures were then combined and incubated at room temperature for 20 min in the dark to prepare the transfection reagent complex. The transfection reagent complex was added to 1×10^6^ HaCaT cells and cultured in a cell incubator containing 5% CO_2_ at 37℃. Simultaneously, a blank control group and a pDsRed-N1 group were set up, with three replicates per group. HaCaT cells from each group were collected, and TvAP65 gene expression was detected by qPCR (primer sequence is shown in Table 2) and western blot (see additional file 4: Figure S4).

### T. vaginalis infection

For *in vitro* infection: HaCaT cells were plated in a 6-well cell culture plate, 1×10^6^ per well. After cells adhered, they were washed twice with pre-warmed PBS, and 1×10^6^ *T. vaginalis* and 2 mL DMEM medium (Procell Life Science & Technology Co., Ltd.) were added per well and cultured in a cell incubator containing 5% CO_2_ at 37℃.

For *in vivo* infection: On the first and eighth days, 500 μg estradiol valerate dissolved in 100 μL sesame oil (Solarbio Science and Technology Co., Ltd., Beijing China) was subcutaneously injected into 6-week-old female BALB/c-nu mice. On the sixth to the ninth day, intraperitoneally inject each mouse with dexamethasone (10 mg/kg, dissolved in 100 μL of PBS) daily for four consecutive days. On day 10, inoculate the mouse vagina with 10 μL of TYM medium (TUOPU Biol-engineering Co., Ltd.) containing 1×10^6^ *T. vaginalis* trophozoite.

### HPV infection rate detection in HaCaT cells

Wild-type *T. vaginalis* or *T. vaginalis* with low TvAP65 expression were infected with wild-type HaCaT cells, TvAP65 overexpressing HaCaT cells, or HaCaT cells with low expression of HSPG2, CD151, or TvAP65 interacting molecules. After 24 h, the medium was sucked away, the cells were washed three times with pre-warmed PBS, and DMEM medium containing RFP-HPV18 pseudovirus particles (Kemei Borui Technology Co., Ltd., Beijing, China) at a final concentration of 5×10^5^ TU/mL was added. After 48 h, the cells were washed three times with pre-warmed PBS, and the infection rate of HPV with red fluorescent markers in HaCaT cells was detected by fluorescence microscopy or flow cytometry.

### HPV infection detection in mouse vaginal tissue

On day one, wild-type *T. vaginalis* or *T. vaginalis* with low TvAP65 expression were infected with 6-week-old female BALB/cA-nu mice. On the second day, 100 μL of 30 mg/mL medroxyprogesterone acetate (Yuanye Bio-Technology Co., Ltd., Shanghai, China) was subcutaneously injected into each mouse. On the sixth day, the mouse vagina was mechanically injured using a sterile cell brush, and 9 μL of RFP-HPV18 pseudovirus particles at a concentration of 5×10^5^ TU/mL were mixed with 3 μL of 4% carboxymethylcellulose (CMC; Yuanye Bio-Technology Co., Ltd., Shanghai, China) and inoculated into the mouse vagina. After 24 h, experimental animals were euthanised by exposure to 60∼70% CO_2_ for 5 min. The mice were dissected and their vaginal tissue was collected, and HPV infection was detected using an *in vivo* imaging system.

### Detection of HPV membrane receptor molecule expression levels

Wild-type *T. vaginalis* or *T. vaginalis* with low TvAP65 expression were used to infect wild-type HaCaT cells, TvAP65 overexpressing HaCaT cells, or HaCaT cells with low expression of TvAP65 interacting molecules. After 24 h, the medium was aspirated and the cells were washed three times with pre-warmed PBS. Total RNA and total protein were extracted. The expression of HPV membrane receptor molecules (HSPG2, CD151, ITGA6, or ITGB1) was detected by qPCR (primer sequence is shown in Table 2) and western blot.

### Statistical analysis

Statistical analysis was performed using SPSS 22.0 software, and graphical analysis was conducted using GraphPad Prism 9.0 software. Image J software was used for greyscale analysis of western blot target bands and in vitro imaging fluorescence intensity. The measurement data were all represented by mean ± standard deviation (x±s). The pairwise comparisons were performed by t test. The comparison of multiple groups was performed by One-way ANOVA and two-way ANOVA. *P* < 0.05 indicated that there was statistical difference between the control group and the experimental group.

## Results

### *T. vaginalis* promotes HPV infection

Fluorescence microscopy, flow cytometry, and *in vivo* imaging system were used to observe the effects of *T. vaginalis* on HPV infection. The results showed that HPV infection rates were significantly higher in both HaCaT cells and the vaginas of BALB/cA-nu mice after *T. vaginalis* infection compared to those not infected with the parasite (Figure 1). This provides direct evidence that *T. vaginalis* promotes HPV infection.

**Figure 1.**
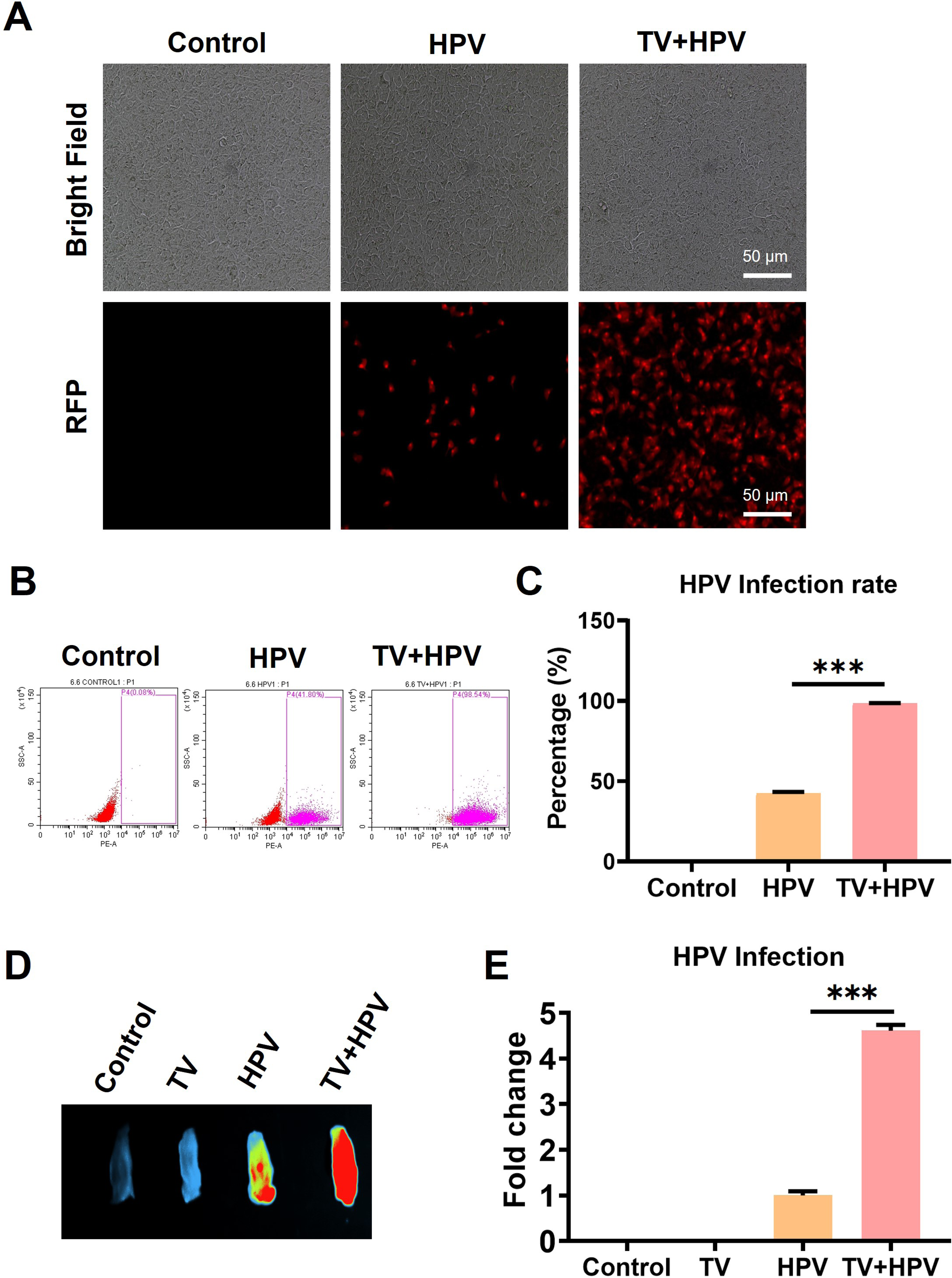
*T. vaginalis* promoted HPV infection. “Control” represents untreated HaCaT cells or mice, “TV” represents mice infected only with *T. vaginalis*, “HPV” represents HaCaT cells or mice infected only with HPV, and “TV+HPV” represents HaCaT cells or mice infected with both *T. vaginalis* and HPV. **A** The HPV infection in HaCaT cells was observed by fluorescence microscopy. **B** The HPV infection rate in HaCat cells was detected by flow cytometry. **C** Flow cytometry data was analyzed using GraphPad Prism 9.0 software. **D** The HPV infection in the vagina of mice was observed by *in vivo* imaging system. **E** The average fluorescence intensity of HPV in the vaginas of mice was analyzed by Image J software. “***” indicates *P*<0.001.

### *T. vaginalis* enhances the expression of HPV membrane receptor molecules

qPCR and western blot were used to detect the effects of *T. vaginalis* on the expression of HPV membrane receptor molecules, including HSPG2, ITGA6, ITGB1, and CD151. The results showed that *T. vaginalis* significantly up-regulated the expression of HSPG2 and CD151, but had no significant effect on ITGA6 and ITGB1 expression levels (Figure 2).

**Figure 2.**
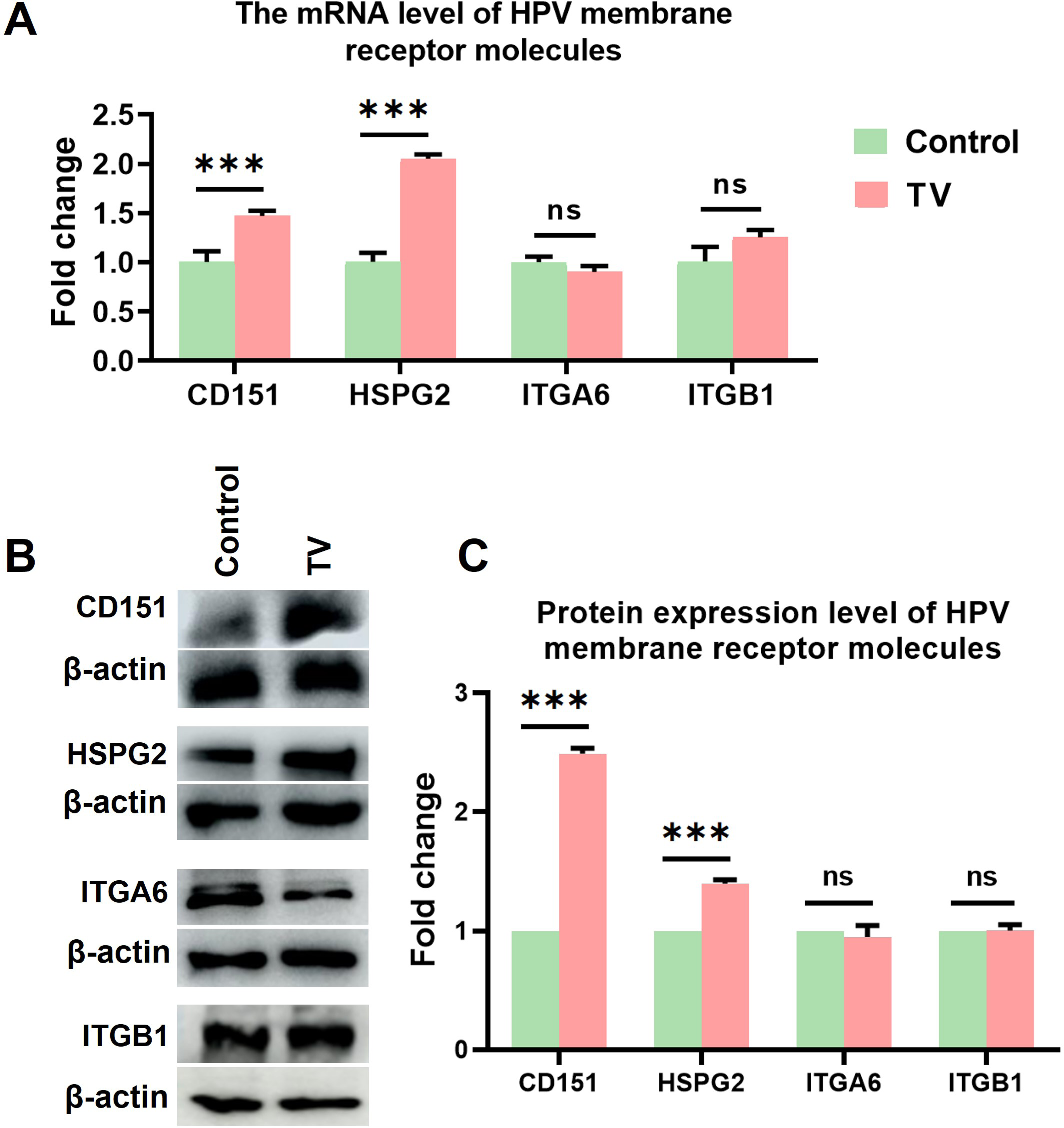
*T. vaginalis* significantly promoted HSPG2 and CD151 expression. “Control” represents HaCaT cells that have not been treated in any way, and “TV” represents HaCaT cells infected with *T. vaginalis*. **A** qPCR was used to detect the mRNA levels of HSPG2, CD151, ITGA6 and ITGB1 in HaCaT cells; **B** WB was used to detect the protein expression of HSPG2, CD151, ITGA6 and ITGB1 in HaCaT cells; **C** Image J software was used to analyze the protein expression levels of HSPG2, CD151, ITGA6 and ITGB1. “ns” indicates no statistical difference, and “***” indicates *P*<0.001.

### TvAP65’s role in *T. vaginalis*-induced promotion of HPV infection and CD151 and HSPG2 expression

Fluorescence microscopy, flow cytometry, and *in vivo* imaging were used to observe HPV infection in HaCaT cells and the vaginas of BALB/c-nu mice after reducing the expression of TvAP65 in *T. vaginalis*. qPCR and western blot were used to assess the effects of TvAP65 on the expression of HSPG2 and CD151 in host cells. The findings revealed that when TvAP65 expression was reduced, the promoting effects of *T. vaginalis* on HPV infection and HSPG2 and CD151 expression were reversed in both HaCaT cells and BALB/c-nu mice (Figures 3 and 4). Furthermore, when a eukaryotic expression vector for TvAP65 was introduced into HaCaT cells, it was observed that the overexpression of TvAP65 significantly promoted HPV infection in HaCaT cells and enhanced HSPG2 and CD151 expression compared to the control group (Figures 5 and 6).

**Figure 3.**
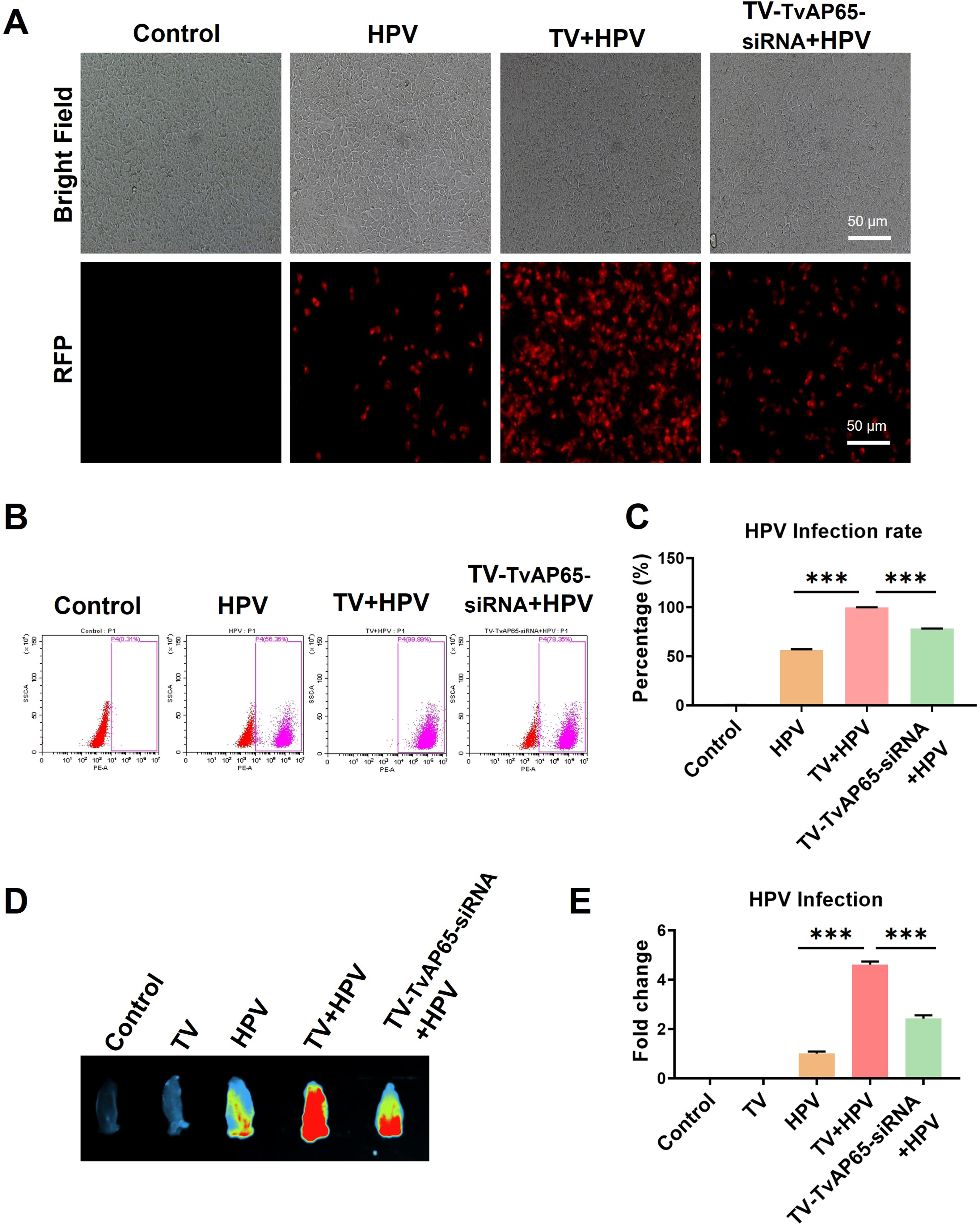
Low-expression of TvAP65 significantly reduced *T. vaginalis*-induced enhancement of HPV infection. “Control” represents untreated HaCaT cells or mice; “TV” represents mice infected only with *T. vaginalis*, “HPV” represents HaCaT cells or mice infected with HPV, “TV+HPV” represents HaCaT cells or mice infected with both *T. vaginalis* and HPV, “TV-TvAP65-siRNA+HPV” represents HaCaT cells or mice infected with both *T. vaginalis* with low-expression TvAP65 and HPV. **A** Fluorescence microscopy observation of HPV infection in HaCaT cells after TvAP65 down-regulation. **B** Flow cytometry detection of HPV infection rates in HaCaT cells after TvAP65 down-regulation. **C** GraphPad Prism 9.0 software analysis of flow cytometry data. **D** *In vivo* imaging system observation of HPV infection in the vagina of mice after TvAP65 down-regulation. **E** Analysis of average fluorescence intensity of HPV in the vaginas of mice using Image J software. “***” indicates *P*<0.001.

**Figure 4.**
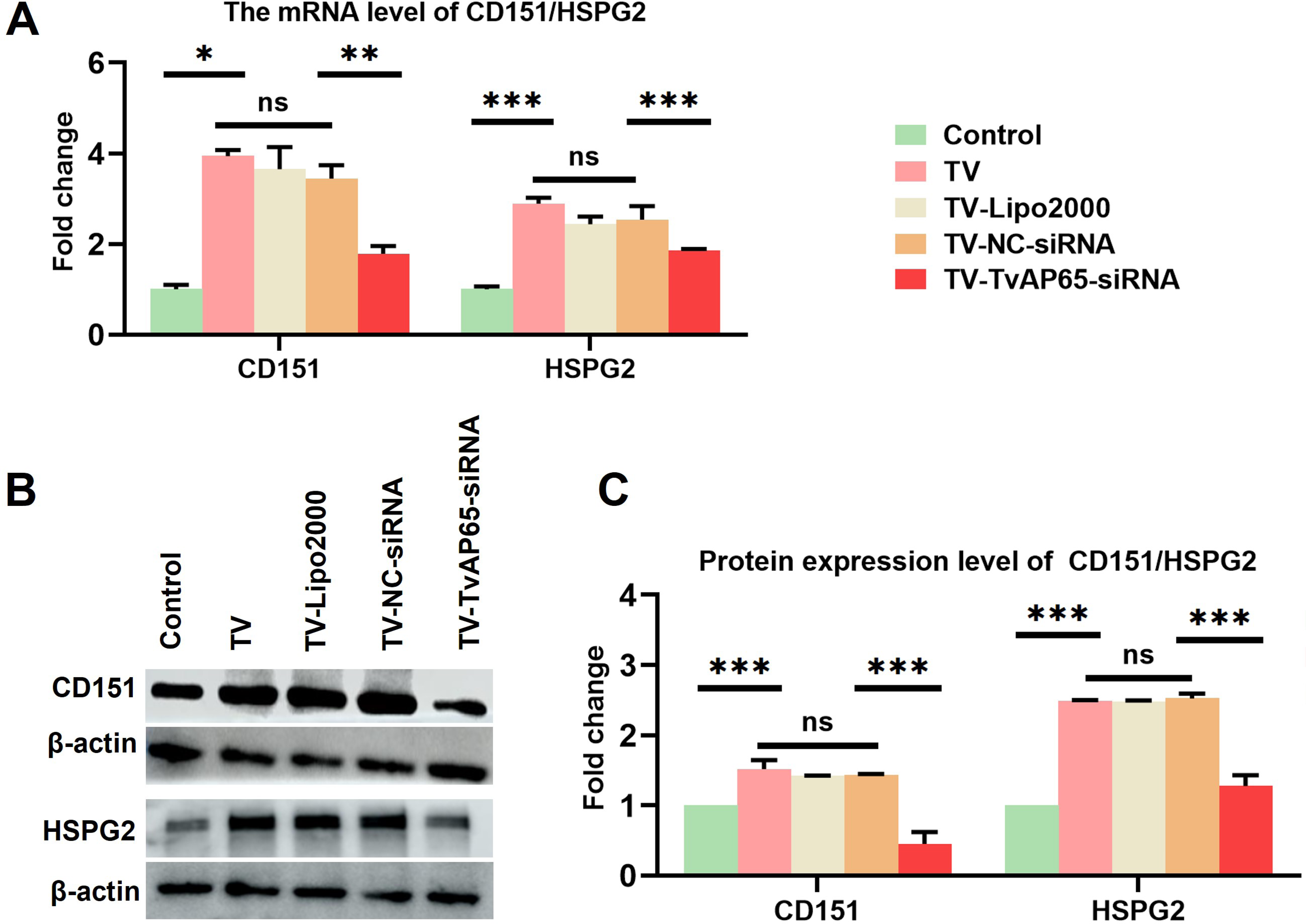
Low-expression of TvAP65 significantly reduced *T. vaginalis*-induced enhancement of CD151 and HSPG2 expression. “Control” represents untreated HaCaT cells, “TV” represents HaCaT cells infected with *T. vaginalis*, and “TV-TVAP65-siRNA” represents HaCaT cells infected with *T. vaginalis* with low TvAP65 expression. “TV-NC-siRNA” and “TV-Lipo2000” served as the control group of siRNA interference. **A** qPCR was used to detect the mRNA levels of HSPG2 and CD151 in HaCaT cells; **B** WB was used to detect the protein expression of HSPG2 and CD151 in HaCaT cells; **C** Image J software was used to analyze the protein expression levels of HSPG2 and CD151. “ns” indicates no statistical difference, “*” indicates *P*<0.05, “**” indicates *P*<0.01, and “***” indicates *P*<0.001.

**Figure 5.**
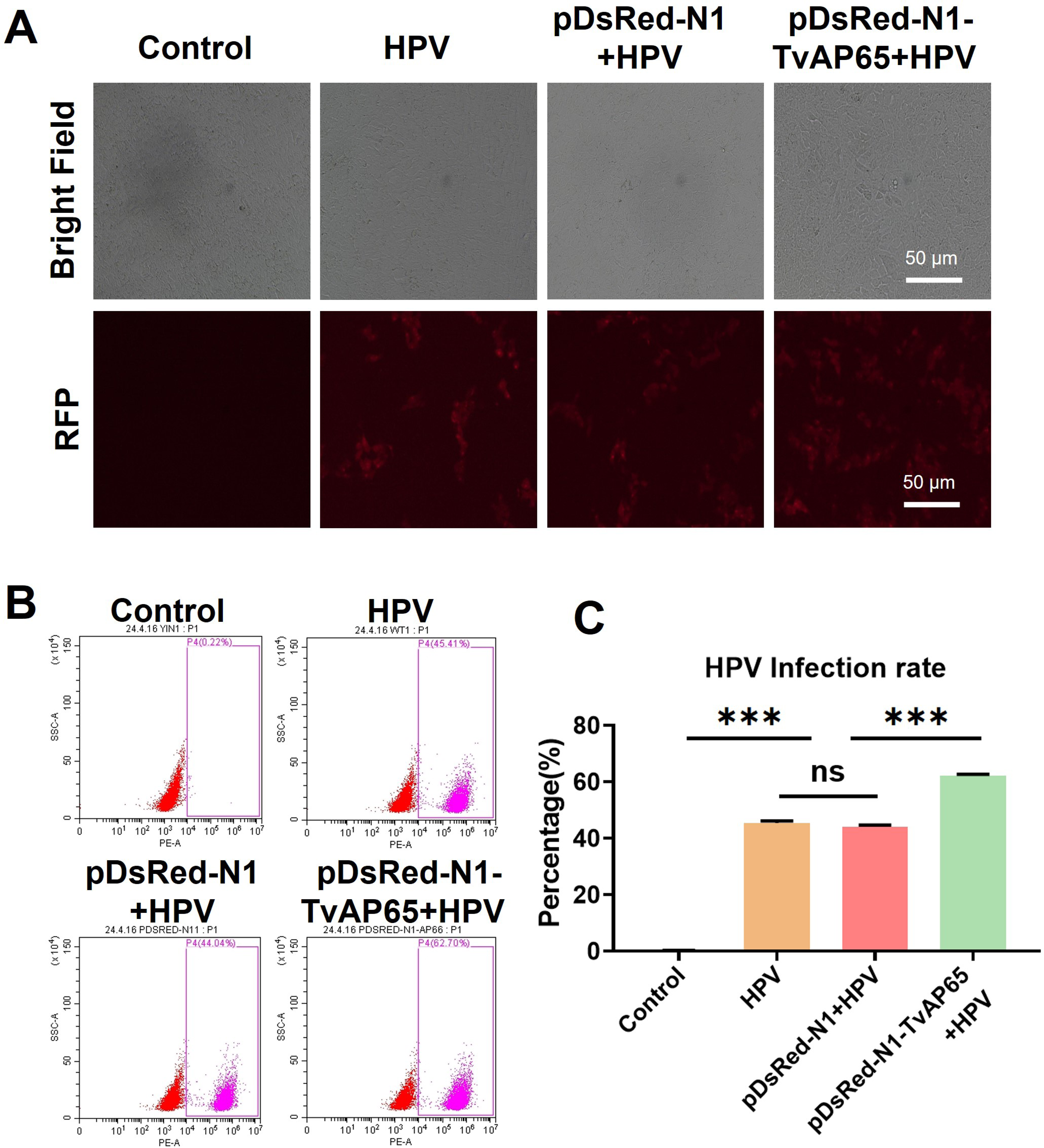
Over-expression of TvAP65 promoted HPV infection of HaCaT cells. “Control” represents untreated HaCaT cells, “HPV” represents HaCaT cells were infected with HPV, “pDsRed-N1-TvAP65+HPV” represents HaCaT cells that over-express TvAP65 were infected with HPV, “pDsRed-N1+HPV” served as a control group of over-expression. **A** The effect of TvAP65 over-expression on HPV infection was observed by fluorescence microscopy; **B** The effect of TvAP65 over-expression on HPV infection was detected by flow cytometry. **C** GraphPad Prism 9.0 software analysis of flow cytometry data. “ns” indicates no statistical difference, and “***” indicates *P*<0.001.

**Figure 6.**
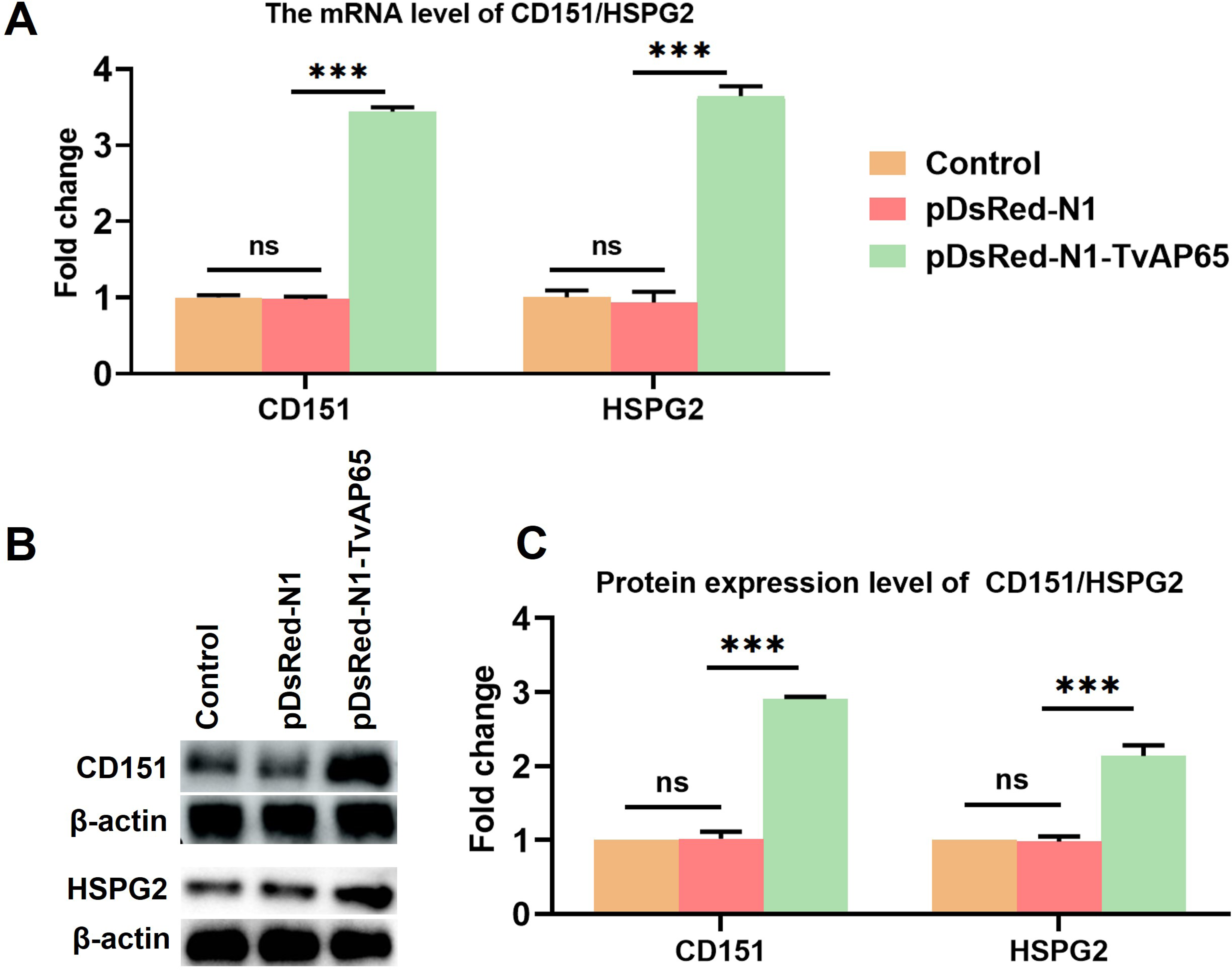
Over-expression of TvAP65 promoted the expression of CD151 and HSPG2 in HaCaT cells. “Control” represents HaCaT cells that have not undergone any treatment, “pDsRed-N1-TvAP65” represents HaCaT cells that over-express TvAP65, “pDsRed-N1” served as a control group of over-expression. **A** The mRNA levels of HSPG2 and CD151 in HaCaT cells were detected by qPCR; **B** The protein expression levels of HSPG2 and CD151 in HaCaT cells were detected by WB. **C** The protein expression levels of HSPG2 and CD151 in HaCaT cells were analyzed by Image J software. “ns” indicates no statistical difference, and “***” indicates *P*<0.001.

### *T. vaginalis* TvAP65 promotes HPV infection by up-regulating CD151 or HSPG2 expression

Fluorescence microscopy and flow cytometry were employed to observe HPV infection in HaCaT cells following the co-downregulation of TvAP65 and HSPG2 or TvAP65 and CD151. It was found that the combined down-regulation of TvAP65 and HSPG2 or TvAP65 and CD151 further reduced the HPV infection rate in HaCaT cells compared to the down-regulation of TvAP65, HSPG2, or CD151 alone (Figure 7). This suggests that *T. vaginalis* TvAP65 promotes HPV infection by up-regulating the expression of CD151 or HSPG2.

**Figure 7.**
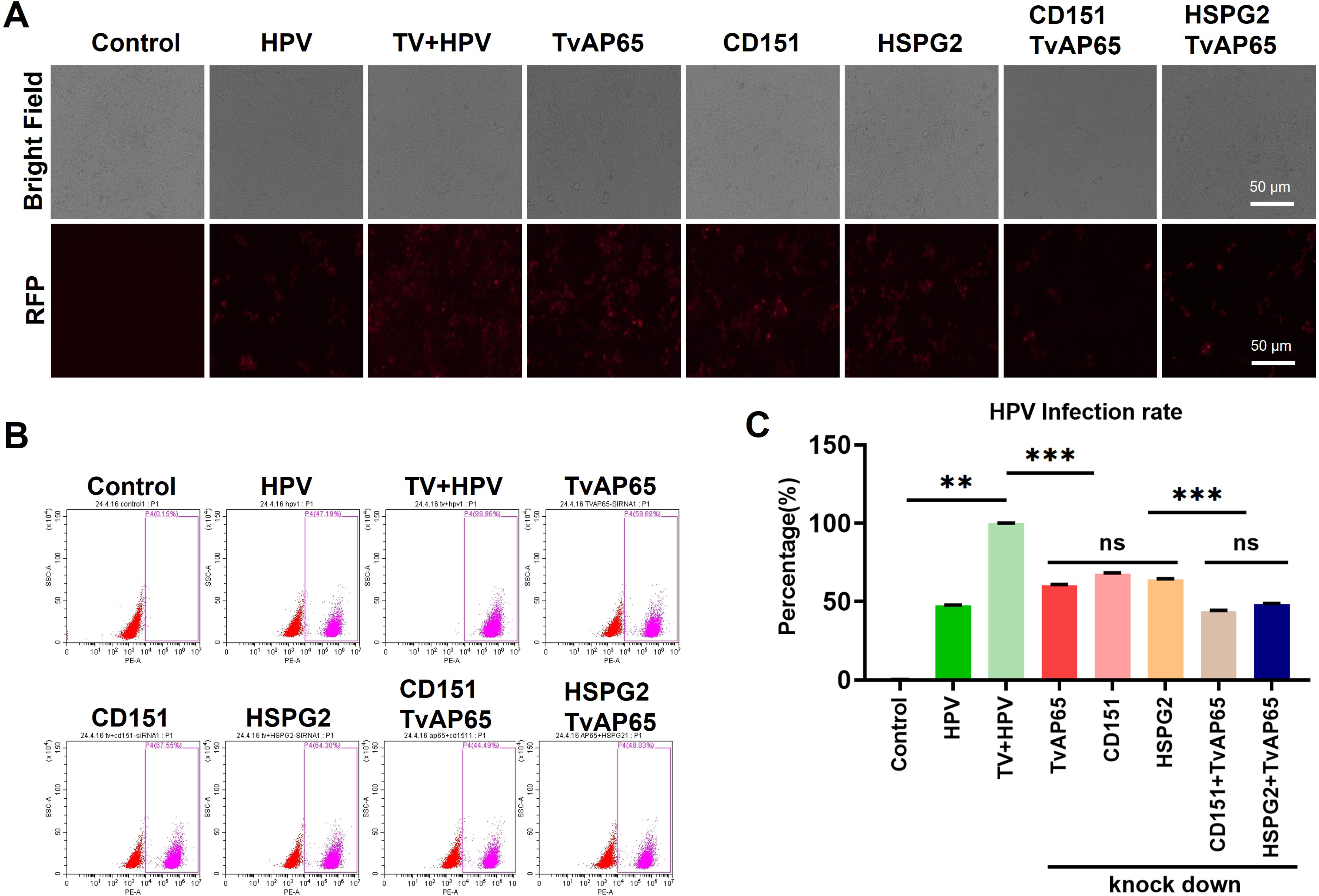
TvAP65 promoted HPV infection by up-regulating CD151/HSPG2 expression. “Control” represents HaCaT cells that have not been treated in any way; “HPV” represents HaCaT cells infected with HPV; “TV+HPV” represents HaCaT cells infected with both *T. vaginalis* and HPV; “TvAP65”, “CD151”, “HSPG2”, “CD151/TvAP65”, and “HSPG2/TvAP65” represent HaCaT cells infected with both *T. vaginalis* and HPV under low-expression of TvAP65, CD151, HSPG2, CD151/TvAP65, or HSPG2/TvAP65 genes, respectively. **A** The HPV infection in HaCaT cells was observed with fluorescence microscope after different genes were down-expressed; **B** and **C** The HPV infection rate in HaCaT cells after different gene down-expression was detected by flow cytometry. “ns” indicates no statistical difference, “**” indicates *P*<0.01, and “***” indicates *P*<0.001.

### The role of TvAP65 interaction molecules in *T. vaginalis*-induced promotion of HPV infection and CD151/HSPG2 expression

Fluorescence microscopy and flow cytometry were used to observe HPV infection in HaCaT cells after down-regulation of TvAP65 interaction molecules. The results showed that the down-regulation of SPCS1, FTH1, ATP5MC3, IGFBP7, REEP5, and PMEPA1 in HaCaT cells significantly reduced the promoting effect of *T. vaginalis* on HPV infection, with the most significant reduction in HPV infection rate observed after down-regulation of SPCS1 (Figure 8). Additionally, SPCS1 also significantly decreased the promoting effect of *T. vaginalis* on HSPG2 and CD151 expression (Figure 9). Compared to the infection of SPCS1-downregulated HaCaT cells with wild-type *T. vaginalis*, the infection rate of HPV significantly decreased further when SPCS1-down-regulated HaCaT cells were infected with TvAP65-down-regulated *T. vaginalis* (Figure 10). This result indicates that the down-regulation of TvAP65 interaction molecules significantly reduces the promoting effect of *T. vaginalis* on HPV infection and the expression of HSPG2 or CD151.

**Figure 8.**
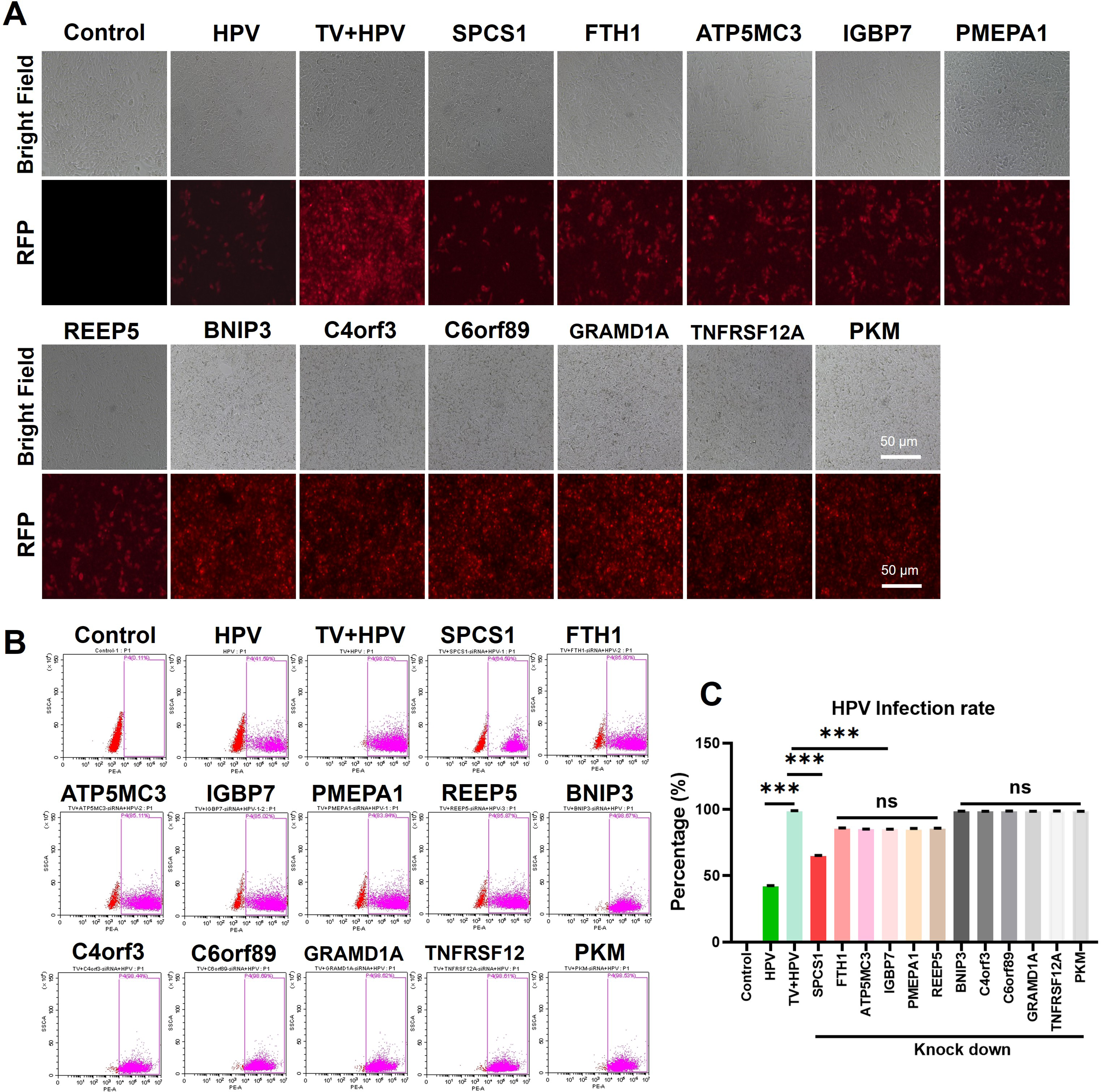
Downregulation of TvAP65 interaction molecule SPCS1 significantly reduced *T. vaginalis*-induced enhancement of HPV infection. “Control” represents HaCaT cells that have not received any treatment; “HPV” represents HaCaT cells infected with HPV; “TV+HPV” represents HaCaT cells infected with both *T. vaginalis* and HPV; “SPCS1”, “FTH1”, “ATP5MC3”, “IGFBP7”, “PMEPA1”, “REEP5”, “BNIP3”, “C4orf3”, “C6orf89”, “GRAMD1A”, “TNFRSF12A”, and “PKM” represent HaCaT cells infected with both *T. vaginalis* and HPV under low-expression of TvAP65 interaction molecules. **A** The HPV infection in HaCaT cells was observed with fluorescence microscope after different genes were down-regulated. **B** and **C** The HPV infection rate in HaCaT cells was detected by flow cytometry after different genes were down-regulated. “ns” indicates no statistical difference, and “***” indicates *P*<0.001.

**Figure 9.**
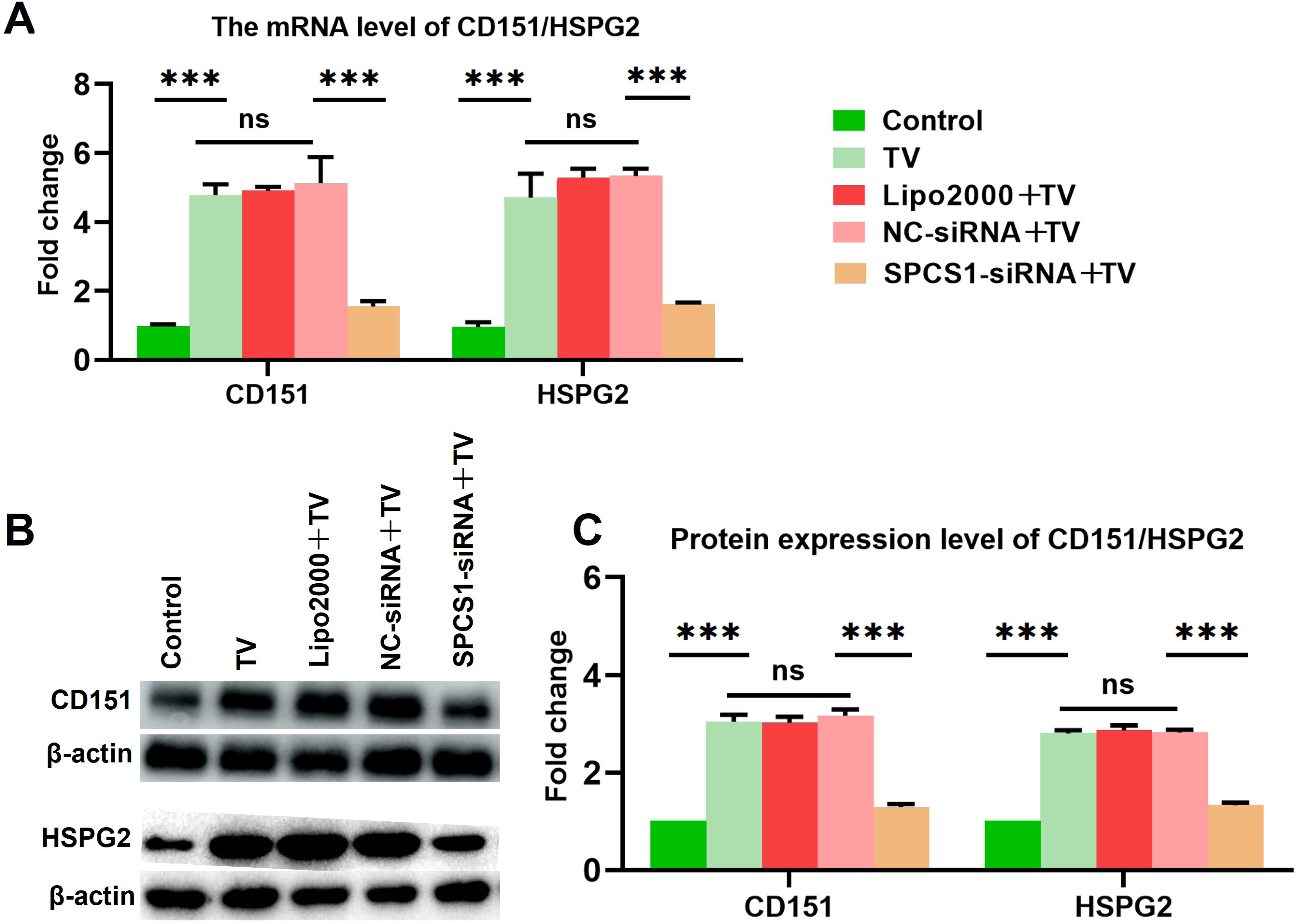
Low-expression of SPCS1 significantly reduced *T. vaginalis*-induced enhancement of CD151 and HSPG2 expression. “Control” represents untreated HaCaT cells, “TV” represents HaCaT cells infected with *T. vaginalis*, and “SPCS1-siRNA+TV” represents that HaCaT cells infected with *T. vaginalis* under low-expression of SPCS1. “NC-siRNA+TV” and “Lipo2000+TV” served as the control group of NC siRNA and Lipofectamine 2000 respectively. **A** qPCR was used to detect the mRNA levels of HSPG2 and CD151 in HaCaT cells; **B** WB was used to detect the protein expression of HSPG2 and CD151 in HaCaT cells. **C** Image J software was used to analyze the protein expression levels of HSPG2 and CD151. “ns” indicates no statistical difference, and “***” indicates *P*<0.001.

**Figure 10.**
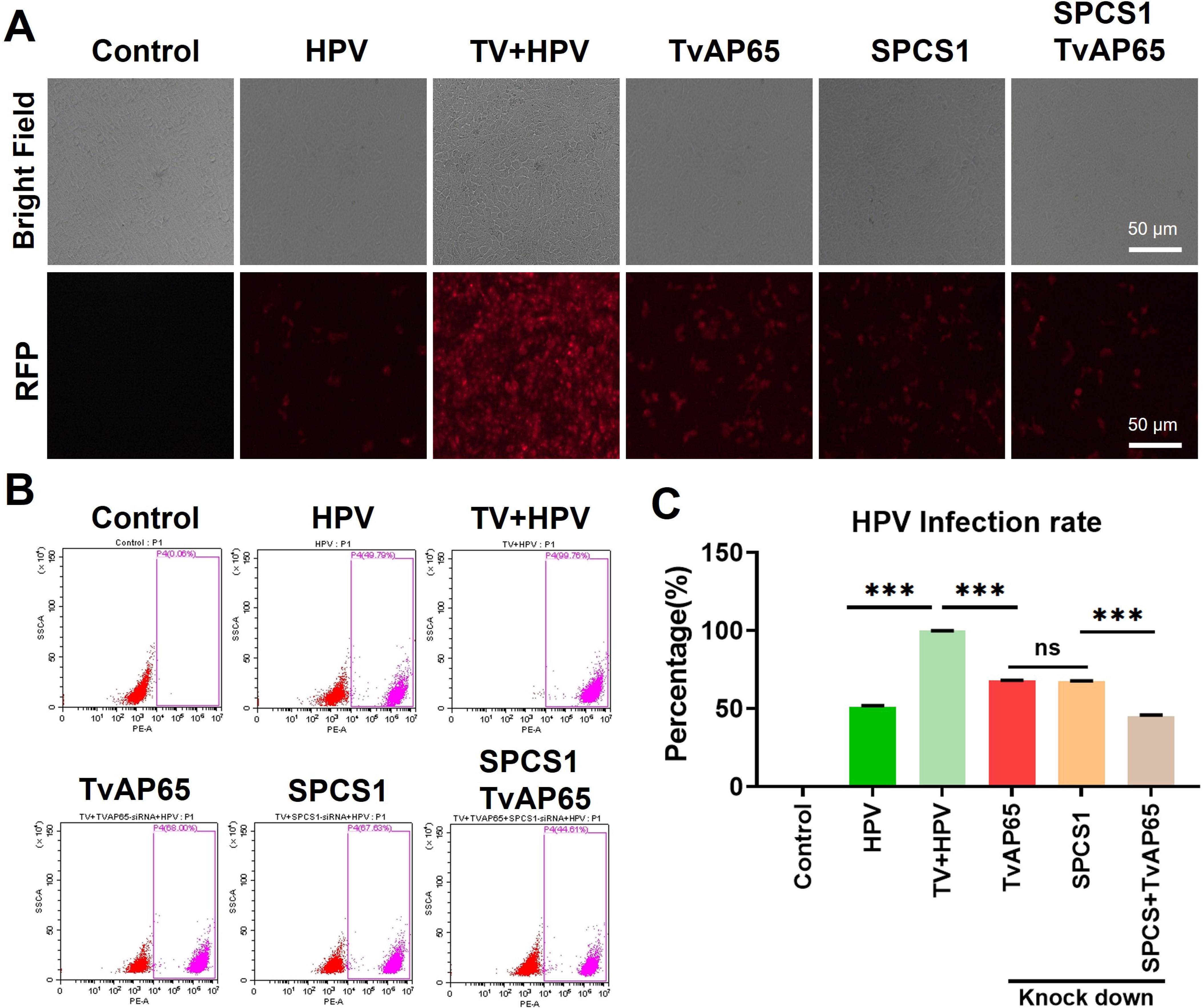
TvAP65 promoted HPV infection by interacting with SPCS1. “Control” represents untreated HaCaT cells; “HPV” represents HaCaT cells infected with HPV; “TV+HPV” represents HaCaT cells infected with both *T. vaginalis* and HPV; “TvAP65”, “SPCS1”, and “SPCS1/TvAP65” represent HaCaT cells infected with both *T. vaginalis* and HPV under low-expression of the TvAP65, SPCS1, or SPCS1/TvAP65 genes, respectively. **A** The HPV infection in HaCaT cells was observed with fluorescence microscopy after low-expression of different genes; **B** and **C** The HPV infection rate in HaCaT cells was detected by flow cytometry after after different genes were down-regulated. “ns” indicates no statistical difference, “**” indicates *P*<0.01, and “***” indicates *P*<0.001.

## Discuss

Sexually transmitted infections (STIs) are major public health issues affecting people worldwide. It is estimated that in 2016, there were 376 million new infections among people aged 15∼49, with an average of more than 1 million cases per day [2]. *T. vaginalis* and HPV are the most common STIs [27]. Multiple epidemiological studies on STIs have pointed out the association between *T. vaginalis* and cervical HPV infection as well as cervical cytological abnormalities [32]. This study further confirmed that *T. vaginalis* can promote HPV infection through *in vivo* and *in vitro* experiments. Further exploration on the interaction mechanism between these two pathogens at the cellular level is of great importance for preventing co-infection with *T. vaginalis* and HPV.

Several studies have reported mechanisms by which different pathogens influence HPV infection. *Gardnerella vaginalis* can promote HPV infection by increasing mucin-degrading enzymes in vaginal secretions, thus compromising the protective mucosal barrier of the host against HPV [33, 34]. *Lactobacillus crispatus* and *Lactobacillus jensenii* can aid in the clearance of HPV infection by inducing immune cells to produce IFN-γ [35]. Regarding parasites, some studies have suggested that substrate production of parasite alters cell membrane and facilitates viral infection. Alternatively, parasites may modulate the cervical microenvironment, thereby facilitating the utilization of vaginal substrates and enhancing pathogen virulence. Additionally, parasites may produce cysteine proteases that degrade antibodies. These pathways can promote the coexistence of parasites and viruses and induce the persistence of pathogens [36, 37, 38]. However, the interaction mechanism between *T. vaginalis* and HPV infection remains unclear.

In this study, we focused on the process of HPV invading host cells and explored how *T. vaginalis* facilitates HPV infection by assisting HPV entry into cells. The HPV invasion process is mediated by multiple cell membrane receptors, including HSPGs, ITGA6, ITGB4, and molecules from the tetraspanin family such as CD63 and CD151 [39]. HPV viruses comprise two capsid proteins, the major capsid protein L1 and the minor capsid protein L2. To trigger the endocytic uptake of viral particles by host cells, the major capsid protein L1 of HPV first binds to the primary membrane receptor HSPGs on the surface of host cells, inducing conformational changes in the virus particles that expose the minor capsid protein L2. Subsequently, the L2 protein binds to ITGA6, the secondary membrane receptor on the surface of host cells, initiating a series of reactions that ultimately result in viral entry [40]. Using inhibitory molecules to block the interaction between HPV and receptors such as HSPG2 or ITGA6 can delay or inhibit HPV entry [41]. Over-expression of ITGA6 can increase the infectivity of HPV pseudovirions to HNSCC cells; blocking ITGA6 with antibodies reduces the binding of HPV to host cells [42]. Our study further discovered that *T. vaginalis* can significantly up-regulate the expression of HPV membrane receptor molecules HSPG2 and CD151 through qPCR and western blot experiments, which maybe an important pathway through which *T. vaginalis* promotes HPV infection.

TvAP65 is not only a major adhesion protein of *T. vaginalis* but also shares sequence identity with the hydrogenosome decarboxylating malic enzyme, indicating that the protein is functionally diverse [43, 44, 45, 46]. Silencing TvAP65 gene can reduce the adhesion of *T. vaginalis* to vaginal epithelial cells without adversely affecting the growth and energy metabolism of *T. vaginalis* [47]. Our study found that the expression level of TvAP65 in *T. vaginalis* was positively correlated with HPV infection rate and HSPG2/CD151 expression level through experiments of low-expression or over-expression of the TvAP65, suggesting that TvAP65 is a key molecule for *T. vaginalis* in promoting HPV infection and up-regulating HSPG2/CD151 expression. To further elucidate whether TvAP65 promoted HPV infection by up-regulating HSPG2/CD151 expression, this study compared HPV infection rates when TvAP65 and HSPG2, or TvAP65 and CD151 were co-silenced with those when TvAP65, HSPG2, or CD151 were silenced alone. It was found that the infection rate of HPV in HaCaT cells was further reduced after co-low expression of TvAP65 and HSPG2 or TvAP65 and CD151, indicating that TvAP65 promotes HPV infection by up-regulating HSPG2/CD151 expression.

When *T. vaginalis* infects host cells, adhesion proteins on the parasite cell membrane bind to specific receptors on the host cell membrane. This interaction forms a unique microenvironment between the parasite and host cell, which is the pathogenic basis for *T. vaginalis* to invade host cells [48, 49, 50, 51]. Studies have shown that TvAP65 interacts with 13 molecules on the host, including FTH1, BNIP3, ATP5MC3, C4orf3, SPCS1, REEP5, PMEPA1, C6orf89, TMEM238, IGFBP7, GRAMD1A, TNFRSF12A, and PKM [31]. Among these, the interaction between TvAP65 and BNIP3 mediates the adhesion and pathogenicity of *T. vaginalis* to host cells. In our study, except for TMEM238 whose sequence had too high GC content to synthesize siRNA, the expression levels of the other 12 molecules in HaCaT cells were individually knocked down. Of these 12 molecules, SPCS1, FTH1, ATP5MC3, IGFBP7, REEP5, and PMEPA1 were involved in promoting HPV infection by *T. vaginalis*, with SPCS1 playing the most significant role. Co-silencing TvAP65 and SPCS1 further significantly reduced HPV expression levels in HaCaT cells. Additionally, this molecule also participates in regulating the expression of HSPG2/CD151 by *T. vaginalis*. These results suggest that TvAP65 in *T. vaginalis* promotes HPV infection by interacting with SPCS1 to up-regulate the expression of HPV membrane receptor molecules HSPG2 and CD151 in HaCaT cells.

In conclusion, *T. vaginalis* promotes HPV entry by up-regulating the expression of cell membrane receptors required for HPV infection (such as HSPG2 and CD151). Particularly, TvAP65 plays a key role in this process, not only enhancing HPV infection rates but also up-regulating the expression of HSPG2 and CD151. Additionally, TvAP65 was involved in the process of promoting HPV infection by interacting with multiple molecules of the host cell, such as SPCS1. Future research could explore the mechanisms of interaction between *T. vaginalis* and HPV at the cellular level, particularly by thoroughly analyzing the specific interactions between TvAP65 and host molecules. This will contribute to comprehensively understand the role of *T. vaginalis* in the process of HPV infection, providing a theoretical foundation for the development of prevention and treatment strategies against co-infection by both pathogens.

## Supporting information

**Additional file 1: Figure S1.** The interference efficiency of TvAP65 siRNAs. **A** Detection of TvAP65 interference effects at the mRNA level by qPCR. **B** Detection of TvAP65 interference effects at the protein expression level by western blot. **C** Gray value analysis of TvAP65 protein expression level using Image J software. “ns” indicates no statistical difference, “***” indicates *P*<0.001.

**Additional file 2: Figure S2.** The interference efficiency of CD151 and HSPG2 siRNAs. **A** Detection of CD151 and HSPG2 interference effects at the mRNA level by qPCR. **B** Detection of CD151 and HSPG2 interference effects at the protein expression level by western blot. **C** Gray value analysis of CD151 and HSPG2 protein expression levels using Image J software. “ns” indicates no statistical difference, “***” indicates *P*<0.001.

**Additional file 3: Figure S3.** The interference efficiency of TvAP65 interaction molecules siRNAs. “ns” indicates no statistical difference, “*” indicates *P*<0.05, “**” indicates *P*<0.01, “***” indicates *P*<0.001.

**Additional file 4: Figure S4**. Detection of TvAP65 over-expression efficiency. A: The mRNA level of the TvAP65 eukaryotic expression vector in HaCaT cells. B: The protein expression of the TvAP65 eukaryotic expression vector in HaCaT cells. C: Analysis of TvAP65 protein expression level using Image J software. “ns” indicates no statistical difference, “***” indicates *P*<0.001.

## Ethical Approval and Consent to participate

The study was reviewed and approved by the Ethics Review Committee of Xinxiang Medical College (Reference No. XYLL-20210184).

## Availability of data and materials

The original contributions presented in the study are included in the article. Further inquiries can be directed to the corresponding author.

## Acknowledgements

Not applicable.

## Consent for publication

Not applicable.

## Author Contributions

XFM, ZCZ and WS designed the study and critically revised the paper. WXS, XFM, WJT, and YNZ prepared the experimental samples and performed the experimental procedures. XFM, WXS and WJT analyzed the results. XFM, ZCZ, WXS, WJT, XWT, ZKY and SW contributed to the writing of the manuscript. All the authors have read and approved the final manuscript.

